# Machine learning approaches identify chemical features for stage-specific antimalarial compounds

**DOI:** 10.1101/2023.08.15.553339

**Authors:** Ashleigh van Heerden, Gemma Turon, Miquel Duran-Frigola, Nelisha Pillay, Lyn-Marié Birkholtz

## Abstract

Efficacy data from diverse chemical libraries, screened against the various stages of the malaria parasite *Plasmodium falciparum*, including asexual blood stage (ABS) parasites and transmissible gametocytes, serves as a valuable reservoir of information on the chemical space of compounds that are either active (or not) against the parasite. We postulated that this data can be mined to define chemical features associated with sole ABS activity and/or those that provide additional life cycle activity profiles like gametocytocidal activity. Additionally, this information could provide chemical features associated with inactive compounds, which could eliminate any future unnecessary screening of similar chemical analogues. Therefore, we aimed to use machine learning to identify the chemical space associated with stage-specific antimalarial activity. We collected data from various chemical libraries that were screened against the asexual (126 374 compounds) and sexual (gametocyte) stages of the parasite (93 941 compounds), calculated the compounds’ molecular fingerprints and trained machine learning models to recognize stage-specific active and inactiv compounds. We were able to build several models that predicts compound activity against ABS and dual-activity against ABS and gametocytes, with Support Vector Machines (SVM) showing superior abilities with high recall (90% and 66%) and low false positive predictions (15% and 1%). This allowed identification of chemical features enriched in active and inactive populations, an important outcome that could be mined for essential chemical features to streamline hit-to-lead optimization strategies of antimalarial candidates. The predictive capabilities of the models held true in diverse chemical spaces, indicating that the ML models are therefore robust and can serve as a prioritization tool to drive and guide phenotypic screening and medicinal chemistry programs.

**For Table of Contents Graphic Only:** 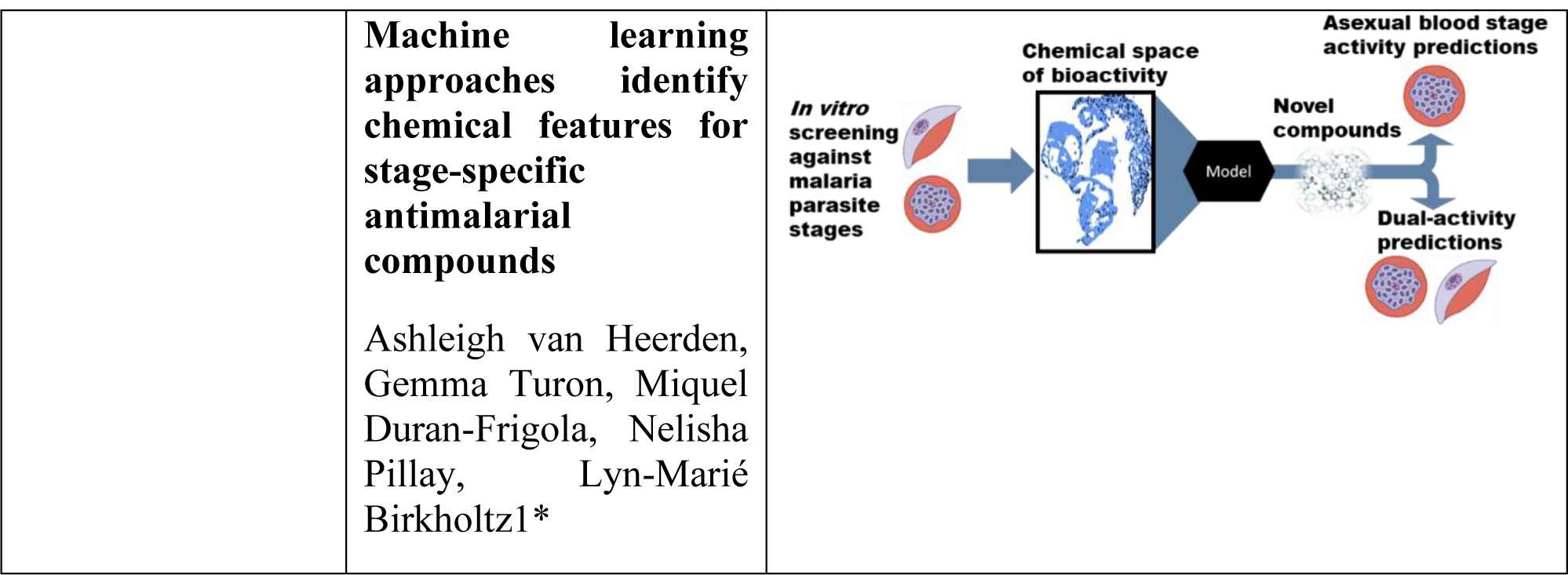

## 1. INTRODUCTION

From 2000 to 2019, malaria-associated deaths showed a steady decline but these gains were stalled in 2020, with a 12% increase in malaria mortality reported globally ^1^. Compounding factors includes the COVID-19 pandemic, which hindered control efforts and the continued emergence of drug-resistant malaria parasites ^2^. Therefore, efforts towards discovering and developing potent antimalarials with novel modes of action must be sustained, and such compounds should target multiple life cycle stages of *Plasmodium* parasites ^3, 4^. Importantly, compounds with the ability to target the transmission of the parasite are sought after as they could be employed to limit the spread of the parasite, and hence disease, and support malaria elimination strategies ^5^. Aside from the required need for compounds with prophylactic activity (blocking exo-erythrocytic development of parasites during liver schizogony), transmission-blocking (TrB) activity can also be ascribed to compounds able to block sexual gametocyte development and subsequent human-to-mosquito transmission of the parasite.

The discovery of compounds with TrB activity through phenotypic, whole-cell screening is fraught with challenges associated with the unique biology inherent to the sexual gametocyte stages of the human malaria parasite, *Plasmodium falciparum*. In this species, only a small portion (∼10%) of the asexual blood stage (ABS) parasites commit to gametocyte development, thereby switching from a proliferative cycle to cellular differentiation. Subsequently, sexually committed parasites differentiate from immature gametocytes (stage I-III) to transmissible, mature stage V gametocytes ^6^. This process is uniquely prolonged amongst the *laverinia* species (*falciparum* and *reichenowi*) and can take ∼10-12 days until mature (stage V) male and female gametocytes are produced as the only forms to support transmission. Identifying gametocytocidal activity is, therefore, more complicated compared to screening for ABS-inhibiting compounds ^5, 7^ due to the limited and time-consuming production of gametocyte populations and the typical need for complementary, orthogonal assays to interrogate the effect of a compound on the differentiated biology associated with transmissible stages ^8^.

Despite these constraints, large chemical libraries have been screened against both the ABS and gametocyte stages of the parasite ^9, 10, 11–14^. This identified compounds with stage-specific activity (e.g., only ABS active) as well as compounds with dual-activity (able to target both ABS and mature gametocytes) or multistage activity (compounds with ABS, TrB and liver-stage activity). These phenotypic screens provide valuable information on the chemical space associated with the differential activity of antiplasmodial compounds. We proposed that this data can be mined to define chemical features associated with compounds displaying activity against only ABS or features in compounds that additionally are active against other life cycle stages including gametocytes. Such information would be very useful to design derivatives of hits during a hit-to-lead medicinal chemistry campaign to enable multistage activity. Additionally, this could exclude compounds that are least likely to show gametocytocidal activity and therefore allow the redistribution of time and resources towards more promising compounds.

Analysing compound activity profiles and relating chemical structural information to bioactivity at scale is an immense and data-heavy task well aligned for machine learning (ML) approaches. The latter have identified important patterns with predictive power in data-dense settings ^15^ and is being increasingly applied to drug discovery ^16, 17^. Whilst proof-of-concept ML models have been generated with fair ability to predict hit compounds with activity against ABS and liver stages independently ^18–20^, this has not been extended to gametocytes. In this study, we expand on and improve these approaches to create an accurate and robust ML classification model able to predict ABS activity and/or gametocytocidal activity, thereby identifying stage-specific as well as dual-active compounds. Our goal was to use more simple models such as SVM and RF rather than complex models such as neural networks. Emphasis was also placed on capturing the chemical space from phenotypic screening data using classical and interpretable molecular fingerprints that allow ML modelling over a broad range of bioactivity prediction tasks ^20–22^.

Since phenotypic screening data is severely imbalanced with few active compounds compared to inactive compounds, we additionally used a hybrid approach to allow the training of models on imbalanced data by combining cluster-based undersampling and algorithms that can implement stronger penalties when misclassifying minority classes. This allowed us to retain relevant chemical space information whilst decreasing the class-imbalance severity to enable model building. We propose that the resultant models will be highly beneficial to accelerate antimalarial drug discovery and development of multistage active antiplasmodial compounds by prioritizing compounds that show the highest probability of desired activity toward a specific stage of the parasite. Not only can this reduce the wasteful expenditure of time and resources on compounds with a low probability of showing activity against different stages of the parasite, but these models can also be mined to identify stage-specific chemical features important for activity against the parasite.

## 2. METHODS

### 2.1. Data acquisition, quality control filtering, and pre-processing of chemical library datasets

Inhibition data of chemical libraries screened against either the asexual and/or gametocyte stages of the parasites were acquired and pre-processed (Figure 1A). These chemical libraries contain multiple chemical spaces, each with several chemically similar analogues. Ultimately, two databases were formed: 1) an ABS database that contained SMILES and inhibition data from 5 chemical libraries screened against *P. falciparum* ABS parasites ^3, 9, 11–13, 23^; and similarly 2) a database (referred to as the dual-active database) that contained SMILES and inhibition data from chemical libraries screened against ABS and any of the gametocyte stages (stage I-V) of *P. falciparum*, with the majority of screening data focussed against stage IV/V gametocytes ^12, 23, 24^ (Table 1). As the chemical libraries were screened by different research groups, using different assay platforms, with different thresholds set to define parasite inhibition and/or gametocytocidal activity, the thresholds specified within the respective screens were used as is to define the binary definition of active/inactive compounds for parasite viability inhibition (Supplementary file S1). Inactive compounds were retained for both databases as these are informative to define the relevant chemical space for bioactivity. Compound SMILES were extracted for all compounds included in the respective databases. Uniform manifold approximation and projection (UMAP) analysis was performed to obtain a projection of the chemical composition of the databases generated with the help of the umap-learn python package version 0.5.3 ^25^.

**Figure 1:**
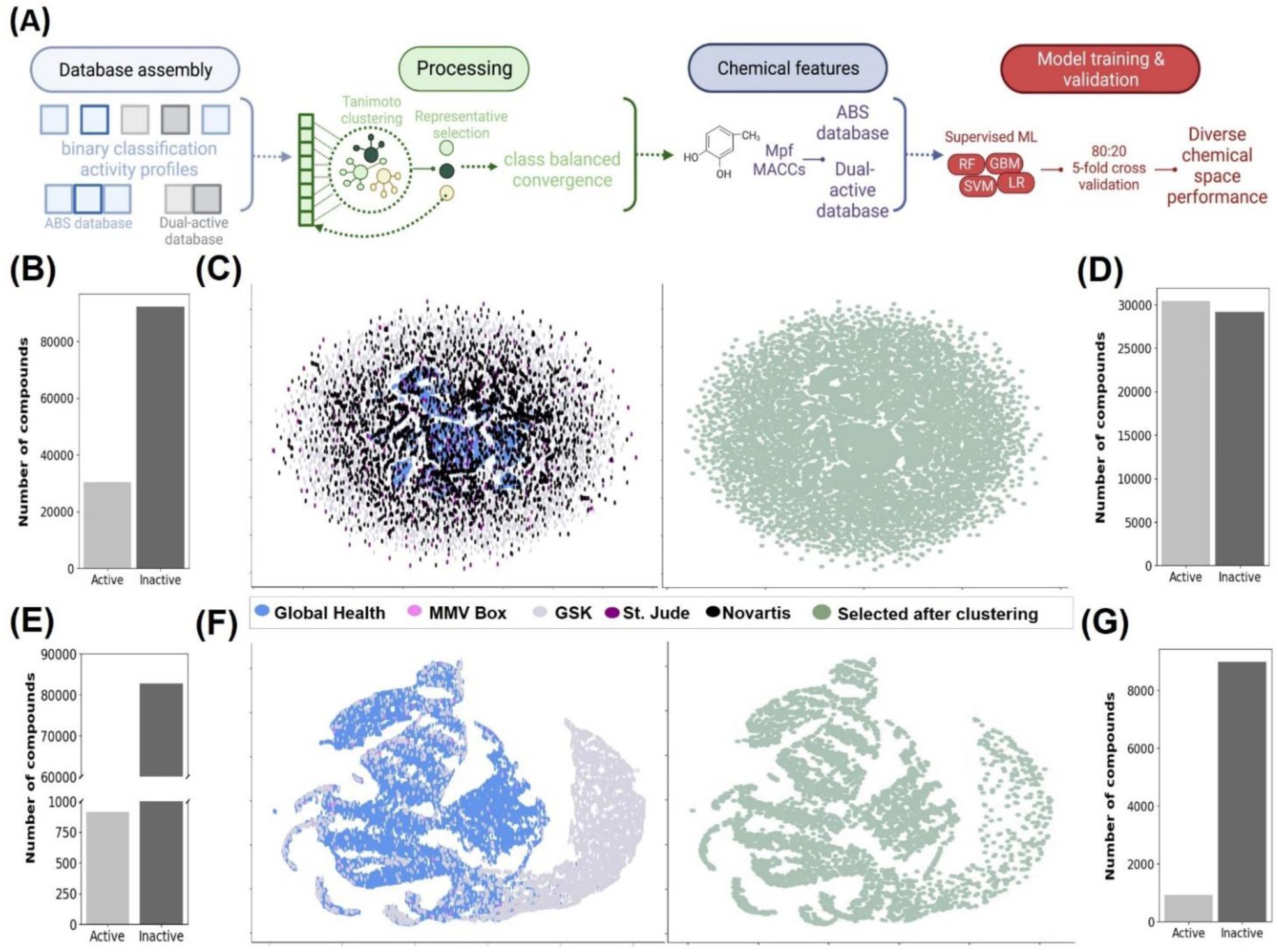
ABS and dual-active database assembly and pre-processing for model building. **(A)** The pipeline used for data assembly, curation, processing, chemical featurization and model building. Data from phenotypic screening of chemical libraries against ABS and/or gametocytes were used for binary definition of active and inactive compounds, class imbalance was addressed via cluster-based undersampling and compounds converted into molecular descriptors Mfp or MACCS to allow model building for compound activity prediction (Created with BioRender.com). **(B** and **E)** Class imbalance in the ABS **(B)** and dual-active **(C)** datasets after binary classification of activity vs inactivity based on criteria as specified in the original screens. **(C** and **F)** UMAP projection of the chemical space in the databases before (lefthand image) and after (righthand image) cluster-based undersampling on inactive compounds for each database. **(D** and **G)** Distribution of active vs inactive compounds for the ABS **(D)** and dual-active **(G)** datasets after cluster-based undersampling.

**Table 1:**
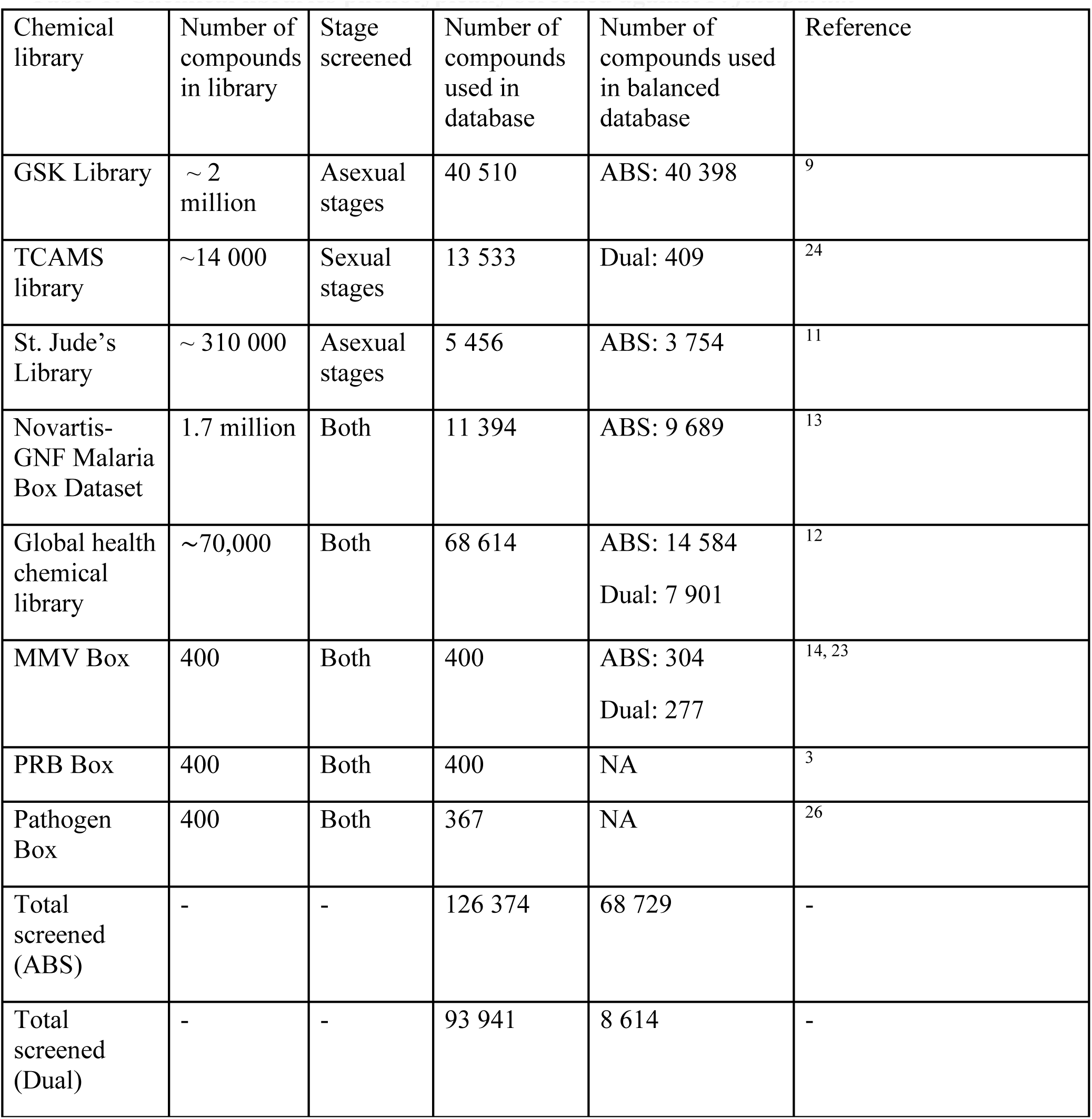
Chemical libraries phenotypically screened against *P. falciparum*.

### 2.2. Cluster-based undersampling of inactive compounds in the databases

The databases created above contained information on both active and inactive compounds on ABS and gametocyte stages of *P. falciparum* parasites. However, the data is inherently skewed towards inactive compounds and this necessitated implementation of class balancing to address the class bias (Figure 1A). We therefore performed cluster-based undersampling using Tanimoto dissimilarity (0.4 distance threshold) with RDKit version 2022.9.5 ^27^ to allow chemical substructure searching and clustering of inactive compounds with similar substructures. The aim was not to completely cluster the chemical databases but to use clustering to aid in the undersampling of inactive compounds. This was independently performed for the ‘ABS’ and ‘dual-active’ databases.

To perform clustering using Tanimoto dissimilarity, compound SMILES of the respective databases were converted RDKit molecular fingerprints (parameter maxpath = 5). Compounds were divided into 7 subsets of 15 – 20 000 compounds each for the ABS database but only 3 subsets of similar size for the dual-active database to reduce the data density and allow the computation of clustering. Parallel clustering was then applied to each of these subsets individually. From this, a representative compound was randomly selected from each inactive cluster within each of the individual subsets. All these identified representatives for the clusters were merged into a single set before chemical clustering was repeated. This process was continued until the database was considered balanced once the number of representatives (inactive compounds) were equal to or >95% of the number of active compounds. For the dual-active database, complete balancing could not be achieved with cluster-based undersampling and the iterative clustering process was halted before reaching the minimum number of clusters to prevent severely limiting the chemical space and causing difficulty in pattern detection in ML.

To determine if cluster-based undersampling allowed better model performance than imbalanced data that had not been pre-processed (undersampled or oversampled), models trained on cluster-based, undersampled data were compared to models trained on imbalanced data. Similarly, to ensure cluster-based undersampling was a better class imbalance correction technique compared to oversampling, the performance of models trained on cluster-based undersampled data, was compared to models trained on oversampled data pre-processed via SMOTE version 0.11.0 ^28^.

### 2.3. Conversion of SMILES to molecular fingerprints for ML

For each of the undersampled databases, canonical compound SMILES were extracted and converted into molecular fingerprints representing the respective compound to allow ML (Figure 1A). Two molecular fingerprints, Morgan fingerprint (Mfp) and MACCS, were generated for each compound to successfully relate structural information for subsequent ML applications ^16,21^. The best bit-length for Mfp was determined to be 500-bits by model evaluation on different bit-lengths (Supplementary Figure S1). Ultimately, a Morgan Fingerprint (Mfp) was generated with RDKit ^27^ that produced a 500-bit feature matrix (5 atoms distance per bit) containing the chemical group and substructure information of the compound. Secondly, MACCS keys were generated for each compound in DeepChem version 2.7.1 ^29^, to generate a 166-feature matrix containing certain chemical groups that have been identified as important within drug discovery and virtual screening ^30^.

### 2.4. Training supervised ML models on the ABS and dual-active database

For each chemical databases (ABS and dual-active), two models were built for each of the algorithms used, one using Mfp as molecular descriptors, the other using MACCS. This was to allow comparative evaluation of the best molecular fingerprint to predict bioactivity in either ABS and/or gametocytes stages in representative and novel chemical spaces.

For model building, the respective databases were randomly split into training and testing sets at an 80:20 ratio, whereby 80% of the compounds is used to train models and the resultant 20% was merged with compounds excluded during undersampling to generate an imbalanced test set for model evaluation on untrained data. Models underwent a grid search cross-validation hyperparameter tuning to identify the optimal hyperparameters (Supplementary Table S1). Each hyperparameter-tuned model then underwent 5-fold cross-validation to assess average accuracy and variability within model predictions.

Since class imbalance was still present (particularly within the dual-active database) and to prevent class bias within models, ML algorithms that applies weight-based mechanisms/penalties on the misclassification of minority classes (dual-active compounds) or other mechanisms for training on imbalanced data were subsequently selected for model building from the scikit-learn python package version 0.20 ^31^. This included ensemble methods such as random forest (RF) and gradient boosting machines (GBM) that have been shown to perform well on imbalanced data ^32^. Additionally, single classifiers such as support vector machines (SVM) and logistic regression (LR) was also applied as these algorithms can attribute weights to a minority class (active compounds), thereby penalising the model more heavily for misclassifying active compounds.

For the balanced ABS database, the ABS activity prediction models were trained on 80% (47 530) of the compounds, whereby the models had to identify patterns within molecular fingerprints of compounds (Mfp or MACCS keys) for the correct prediction of compound bioactivity against ABS. For SVM, RF, GBM and LR models the scikit-learn python package was used to build and train the model to training set. During training, ABS activity models were built using the optimal hyperparameters identified (Supplementary Table S1) and underwent subsequent 5-fold cross-validation. Thereafter, the imbalanced test set (61 029) excluded from training, was used to assess the model bioactivity prediction accuracy and overfitting on imbalanced untrained data.

Dual-activity prediction models were similarly trained on 80% (7 913 compounds) of the dual-active database. Similar to the models for the ABS data, scikit-learn was used to build RF and GBM models. For LR models, however, the class weight was additionally set as 1 for inactive compounds and 10 for active compounds to compensate for class imbalance in dual-activity prediction models when training on the training set. Similarly, for SVM the class weight set to “balanced” to adjust class weights inversely proportionally to the frequency of the class. Each dual-activity prediction model was built using the optimal hyperparameters identified (Supplementary Table S1) and subsequently underwent 5-fold cross-validation before evaluation on the imbalanced test set (62 375 compounds) to assess model bioactivity prediction accuracy and overfitting.

### 2.5. Metrics for evaluating different ML algorithms in predicting asexual and dual-activity

Tuned models were assessed on their cross-validation results as well as their test set results to determine the model performance on untrained imbalanced chemical data and to highlight any overfitting within the models. Metrics used for model predictions evaluation were recall (equation A), precision (B), false positive rate (C), receiver operator characteristic curve (ROC-AUC) and the F1-score (D).

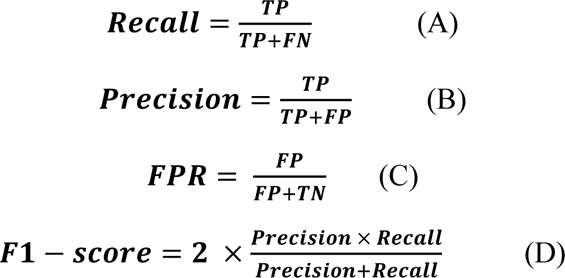

Recall and precision of models was calculated to determine whether models were able to correctly predict active and inactive compounds whereas FPR assess false positive predictions. The ROC-AUC score was used to evaluate how well the model can distinguish between two classes. A score of 0.7 indicates the model has a 70% chance to correctly distinguish active and inactive compounds. In addition, we also used the F1-score, which combines recall and precision^33^. Besides precision and recall, models were additionally evaluated for optimised sensitivity (equation E below) and specificity (F), used to define the geometric mean (G), when predicting active and inactive compounds within novel chemical spaces.

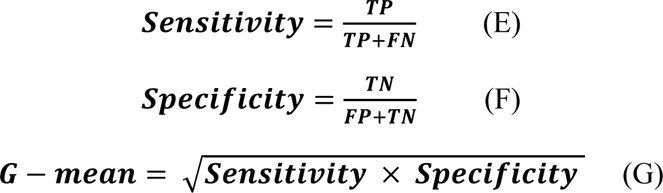

Considering the class imbalance present, especially within the dual-active database, the GHOST python package version 0.6.1 ^34^ was used to adjust the probability thresholds for model decision and to examine if this enabled better model performance upon the test set to the metrics mentioned above. To determine whether such simplistic models is on par or even better than complex models such as neural networks, the top performing model based on these metrics would then be compared to more complex models generated via the autogluon python package version 0.8.2 ^35^ using the same training and test set data.

### 2.6. Chemical features important for predicting ABS inhibition and dual-activity

These compound activity prediction models hold important chemical space information for activity prediction. The aim was to use the models to identify chemical features that are both statistically significant and important for the model in compound activity prediction. Some models have the ability to rank features (i.e. chemical features) according to the weight/importance such features have in predicting activity or inactivity of a compound. Unfortunately, highlighting the Mfp enriched features important for predicting dual-activity was complicated due to our best performing model, SVM, employed a radial basis function (RBF) kernel transformation technique which complicated how one can calculate the true weight a feature holds within model predictions. To allow interpretability RF models trained on Mfp with good precision and ability to distinguish active and inactive compounds within representative chemical spaces against ABS and/or gametocytes, were selected to perform feature importance analysis. To highlight such features important for activity prediction, Mfp features significantly enriched within active compounds (ABS inhibiting or dual-active) were identified using the Z-test on two sample proportions. From the Z-scores calculated using Z-test, Mfp features significantly enriched (Z-score >2.5, p-value < 0.01) within active compounds compared to inactive compounds were identified and then ranked according to the feature importance score obtained from the best performing activity prediction model. From this the top 100 Mfp features were identified and compared to the top 100 Mfp features of other activity prediction models to highlight any overlap between the top Mfp features of different models. This assumes that if features are predictive of compound activity, such features should overlap as the top features for different models that employ different algorithms in pattern recognition.

### 2.7. External validation of ML performance on chemically diverse PRB and Pathogen Box

To evaluate the limits of activity prediction for the models in novel chemical spaces, and to externally validate the models, models were additionally tested against new chemical libraries that included chemically diverse compounds. This included the data from the open-source Medicines for Malaria Venture (MMV) Pandemic Response Box (PRB box) and the Pathogen Box, with potent activity against various stages of the parasite (www.mmv.org) ^3, 26^. The hit rate of the best performing model was then compared to that of the chemical library and random selection. Here the hit rate of the top 100 compounds (ordered according to model probability) was compared to the randomly selected 100 compounds. To assess whether the models aided in limiting the number of active compounds in the bottom 100, i.e., least likely to have activity or last to be screened via random selection, the hit rate for the bottom 100 was also calculated with the goal of having a lower hit rate than that of randomly screening. The idea being the top 100 would be the first compounds that are randomly select to start with during phenotypic screening whereas the bottom being the last compounds chosen to screen.

## 3. RESULTS

### 3.1. Database generation and class imbalance correction

A data analysis pipeline was devised to allow database generation, class imbalance correction and processing, chemical featurization and model training and validation (Figure 1A). This pipeline was applied in parallel to create two different modelling environments: 1) an environment for compounds with predicted stage-specific activity against ABS parasites alone, and 2) for compounds with dual-activity against both ABS parasites and gametocytes. Therefore, two databases were generated from datasets of chemical libraries screened against either the asexual and/or gametocyte stages of *P. falciparum* parasites (Figure 1A, Table 1) ^3, 9, 11–13, 23, 24^. The first database included compounds which were screened against ABS and the second database where compounds screened against both ABS and gametocyte stages. Of all active compounds, 96% were those inhibiting ABS and 3% showing dual-activity, with only 0.3% displaying sole activity against gametocyte stages. Binary classifications of activity were retained as per criteria defined within each screen.

Within these datasets, the ratio of inactive to active compounds is inherently skewed towards the inactive compounds, creating imbalanced datasets with inactive compounds comprising 75% of the ABS database compared to a more severe situation of 99% in the dual-active database (Figure 1B and E). To prevent loss of relevant chemical space information present in inactive samples ^36^, cluster-based undersampling was performed on inactive compounds for each database (Figure 1A) to generate balanced datasets more amenable for conventional ML modelling ^37^. This clustering approach could be justified as it was observed that these chemical libraries contained multiple structurally-related compounds within specific chemical spaces that have similar inactivity, which could function as representative compounds (Figure 1C). UMAP analysis and spatial projection of the ABS database indicated retention of the chemical composition and diversity after two rounds of subset clustering of inactive compounds (Figure 1C), with class balancing attained (30 939 active compounds *vs*. 29 143 inactive compounds, Figure 1D). However, after multiple rounds of clustering for the dual-active database, only a maximal of 3 745 clusters representing the inactive compounds were obtained, which did not correct the class imbalance. Therefore, additional parallel chemical clustering was performed, and this process halted before reaching the maximal number of chemical clusters. This resulted in more than one chemical representative of clusters present (8 975 clusters) to define the chemical space for ML for the inactive compounds and corrected the class imbalance to <8-fold (Figure 1E and G). The chemical distribution and composition for inactive compounds (Figure 1F) was also retained for the dual-active database. From these more balanced databases, the chemical clusters were shuffled and randomly split where 80% of compounds were used for model training and the remaining 20% was merged with inactive compounds excluded during cluster-based undersampling to create an imbalanced test set for model evaluation.

To determine if training on cluster-based undersampled data as above skewed model performance on the imbalanced test set, we compared this strategy to oversampling or training on imbalanced data without pre-processing. The models trained on undersampled data for both ABS and dual activity resulted in higher sensitivity but similar specificity compared to models trained on oversampled or severely imbalanced data, with higher G-mean and ROC-AUC scores for the undersampled data (Supplementary file S1). This indicates that the models trained on oversampled data fails to identify active compounds due to the model possibly fixating on patterns associated with the oversampled active compounds in the training data ^38^. Cluster-based undersampling therefore provided a better class imbalance correction technique and improved model sensitivity.

### 3.2. ABS activity prediction model selection and performance

To associate chemical features of compounds with activity against ABS (or lack thereof) during model training, a two-pronged approach was used: the SMILES for compounds from the ABS database was converted into either Mfp or MACCS molecular fingerprints. Subsequently, data from both of these featurization methods were used and four different models each were trained on 80% of the data in each instance (Figure 1A). For the models trained on Mfps of compounds from the ABS database, the SVM model achieved the highest ROC-AUC score with lowest variability (0.99 ± 0.02) during 5-fold cross-validation (Figure 2A). The SVM model maintained similar accuracy when predicting compound ABS inhibition activity on untrained imbalanced test data (ROC-AUC score 0.92, Figure 2B), indicating no overfitting of the model to training data.

**Figure 2:**
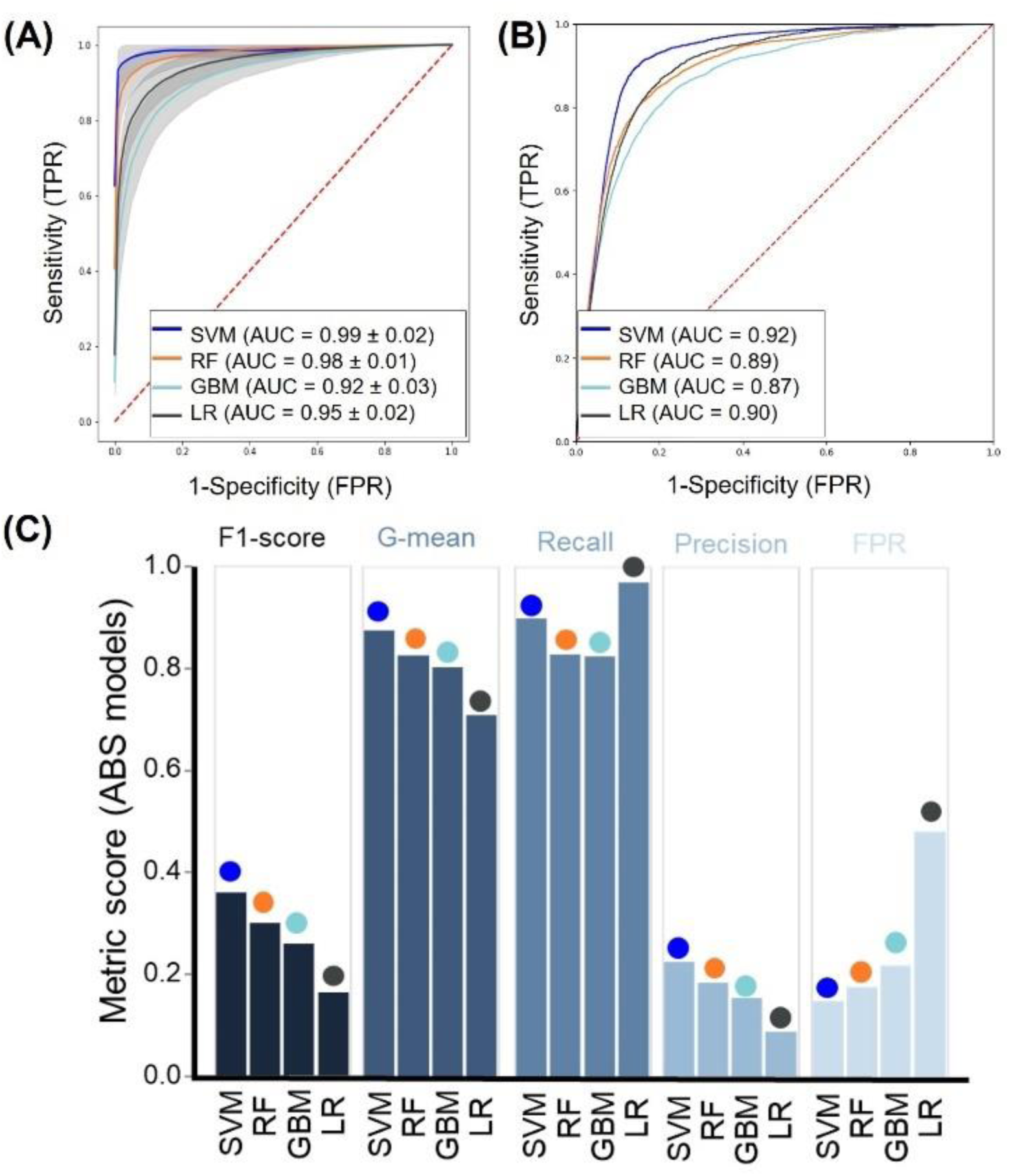
Performance of different ML algorithms in identifying compounds with ABS activity. **(A)** ROC-AUC curves showing performance of different ML algorithms in predicting compounds with ABS activity when trained on the Mfp of compounds after 5-fold cross validation. Insert indicates AUC mean values ± standard deviation. **(B)** ROC-AUC curves showing performance of different ML algorithms on the imbalanced test set. **(C)** Model performance metrics associated with the performance of the different models in predicting the imbalanced test set data. The F1-score evaluated model performance on imbalanced data, whereas G-mean scores determined how well models were able to optimise sensitivity and specificity. Recall and precision indicated accuracy of activity predictions whereas false positive rate (FPR) indicated error within predictions.

Additionally, the SVM models’ recall ability (at 0.90) and precision (at 0.22) was comparable to that of the other ensemble models (RF, GMB; Figure 2B), with only LR models obtaining a higher recall (0.97) than SVM. However, it does this at the expense of the LR model’s precision (0.08), resulting in low specificity with a higher false positive rate (FPR: 0.43 vs <0.22 for the other models) (Figure 2B and 2C). This indicated that the model derived from SVM for ABS activity prediction is more precise and better able to identify compounds with ABS activity whilst limiting its’ false positive rate (FPR: 0.15) (Figure 2C), with losses in FPR associated with gains in precision to as high as 0.675 at higher probability thresholds (Supplementary Table S2). Interestingly, SVM also showed similar, if not better, performance than that of more complex models such as NeuralNetFastAI (Supplementary Table S3) and did much better at reducing its FPR compared to such complex models.

Similar to the data on models trained using Mfps, SVM models trained with MACCS keys of compounds also achieved the highest ROC-AUC scores on both 5-fold cross-validation (0.99 ± 0.02) (Supplementary Figure S2) and untrained test data (0.92, Supplementary Figure S2). No major differences could be detected between the performance of models trained on Mfps or MACCS keys beyond a slight improvement in performance metrics for LR and RF models (Supplementary file S1).

### 3.3. Model selection and performance of dual-activity prediction models

To evaluate the performance of the models to predict compounds with dual-activity, different metrics were used to ensure accurate evaluation of models within the constraints of the class imbalanced dual-active database. Models were evaluated on their recall and precision in identifying dual-active compounds. SVM outperformed other models within 5-fold cross-validation by obtaining ROC-AUC means >0.96 (Figure 3A, Supplementary Figure S2). This extended to the performance of the models against imbalanced test data, where SVM reached ROC-AUC score of 0.95 for models trained on Mfp (Figure 3B, Supplementary Figure S2) indicating no overfitting of models. Considering the large disparity in the number of actives *vs*. inactive compounds for dual-activity, the F1, G-mean and recall scores were also considered, with the SVM models identified as the best performing model trained on Mfp for dual-activity prediction with optimal F1 (0.26), G-mean (0.81), and recall (0.66) scores and low false positive predictions (0.006) (Figure 3C). Higher probability thresholds again resulted in increased precision to 0.562 (Supplementary Table S2) Interestingly, MACCS generally resulted in a higher FPR than models trained on Mfp, indicating that MACCS is not as good at generating molecular descriptors of the chemical space for dual-activity, whereas Mfp in comparison is more extensive and descriptive (Supplementary file S1).

**Figure 3:**
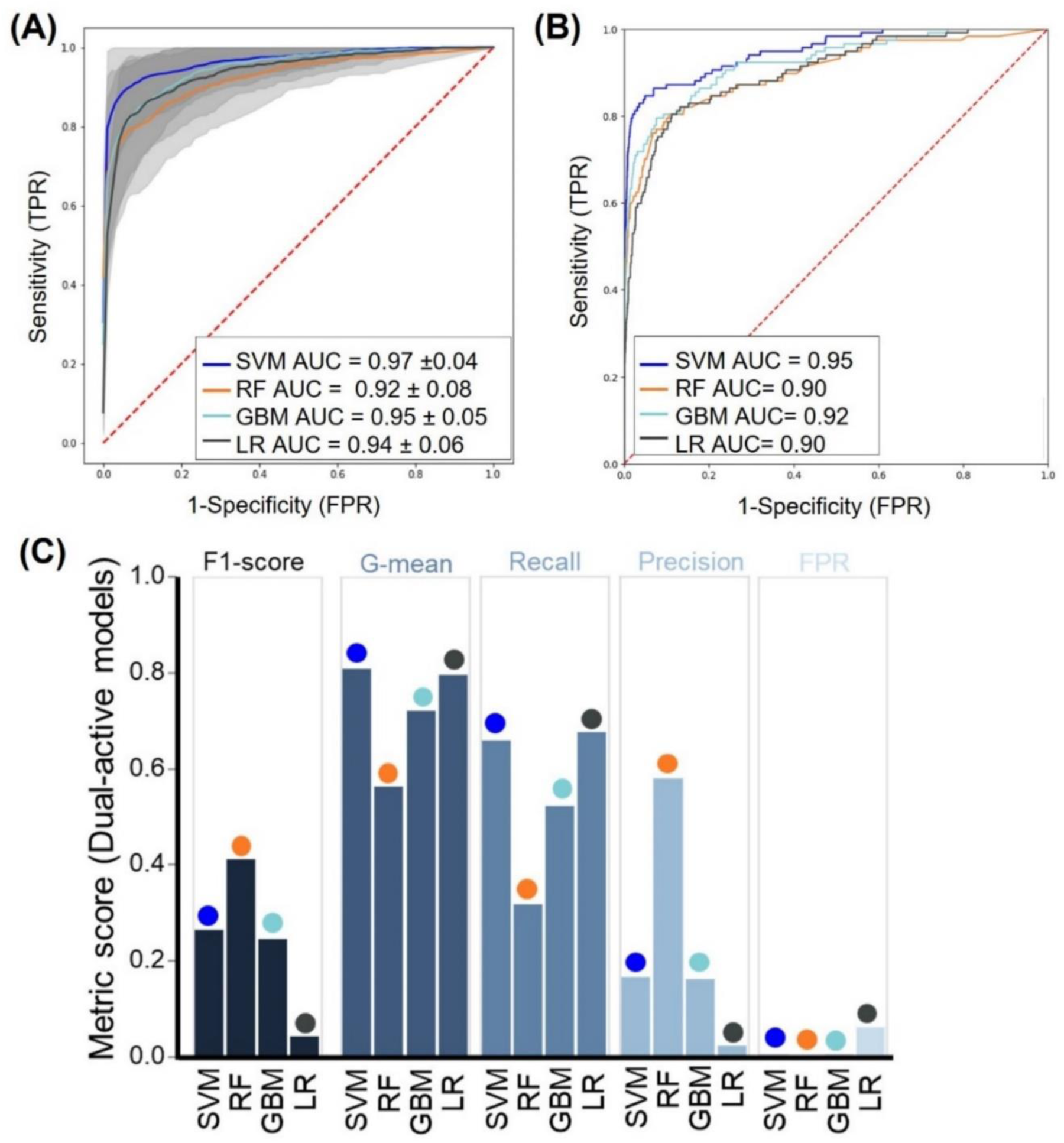
Performance of different ML algorithms in identifying compounds with dual-activity. **(A)** ROC-AUC curves showing 5-fold cross validation performance of different ML algorithms in predicting compounds with dual-activity when trained on the Mfp of compounds. Insert indicates AUC mean values ± standard deviation. **(B)** ROC-AUC curves showing performance of different ML algorithms on the imbalanced test set. **(C)** Model performance metrics associated with the performance of the different models in predicting imbalanced test set data. The F1-score evaluated model performance on imbalanced data, whereas G-mean scores determined how well models were able to optimise sensitivity and specificity. Recall and precision indicated accuracy of activity predictions whereas false positive rate (FPR) indicated error within predictions.

The high ROC-AUC and recall scores of the SVM models were on par with those obtained when more complex models were generated on the same datasets (Table S3). However, we observed that our simple models tended to outperform these more complex models with the SVM models having the lowest FPR, indicating that complex models tend to overfit the data and have poor generalization on these rather small datasets. In conclusion, SVM was the best model for predicting compounds with dual-activity with low false positive predictions on test data within representative chemical spaces.

### 3.4. Identification of top chemical features for activity prediction

Considering the performance of models trained on Mfp as molecular descriptor of chemical compounds’ structural features, it was of interest to determine if there were any features that are particularly associated with antiplasmodial activity, or lack thereof. However, SVM models could not be used for feature importance analysis due to these models using a RBF data transformation technique which complicates attributing importance scores to chemical features. Therefore, RF was used because it was the second-best model in each instance and these models could provide a feature importance score for particular chemical features. This indicated that a larger number of Mfp features were enriched within compounds with ABS activity (383) compared to inactive compounds (67) (Figure 4A). Recursive feature elimination tended to select for Mfp features enriched within inactive compounds (data not shown) and therefore an arbitrary selection of the top 100 features based on model importance scores were applied to provide a more comprehensive, inclusive dataset (Supplementary file S2). More than half of these features identified with the RF models were additionally confirmed by at least one other model. Importantly, there was no overlap between the top 100 features enriched for ABS activity and the features enriched in inactive compounds (Figure 4A, Supplementary file S2), validating that a high level of specificity was obtained when associating Mfp features with ABS activity. For the compounds with dual-activity (Figure 4B), less Mfp features were enriched within active compounds compared to those observed in the ABS space (266 features compared to 383 for ABS activity). However, as with the features describing ABS activity, the features enriched in compounds with dual-activity were specific, with no overlap present between the top 100 Mfp features for dual-activity compared to the 52 features enriched in compounds that are inactive against gametocytes (Figure 4B).

**Figure 4:**
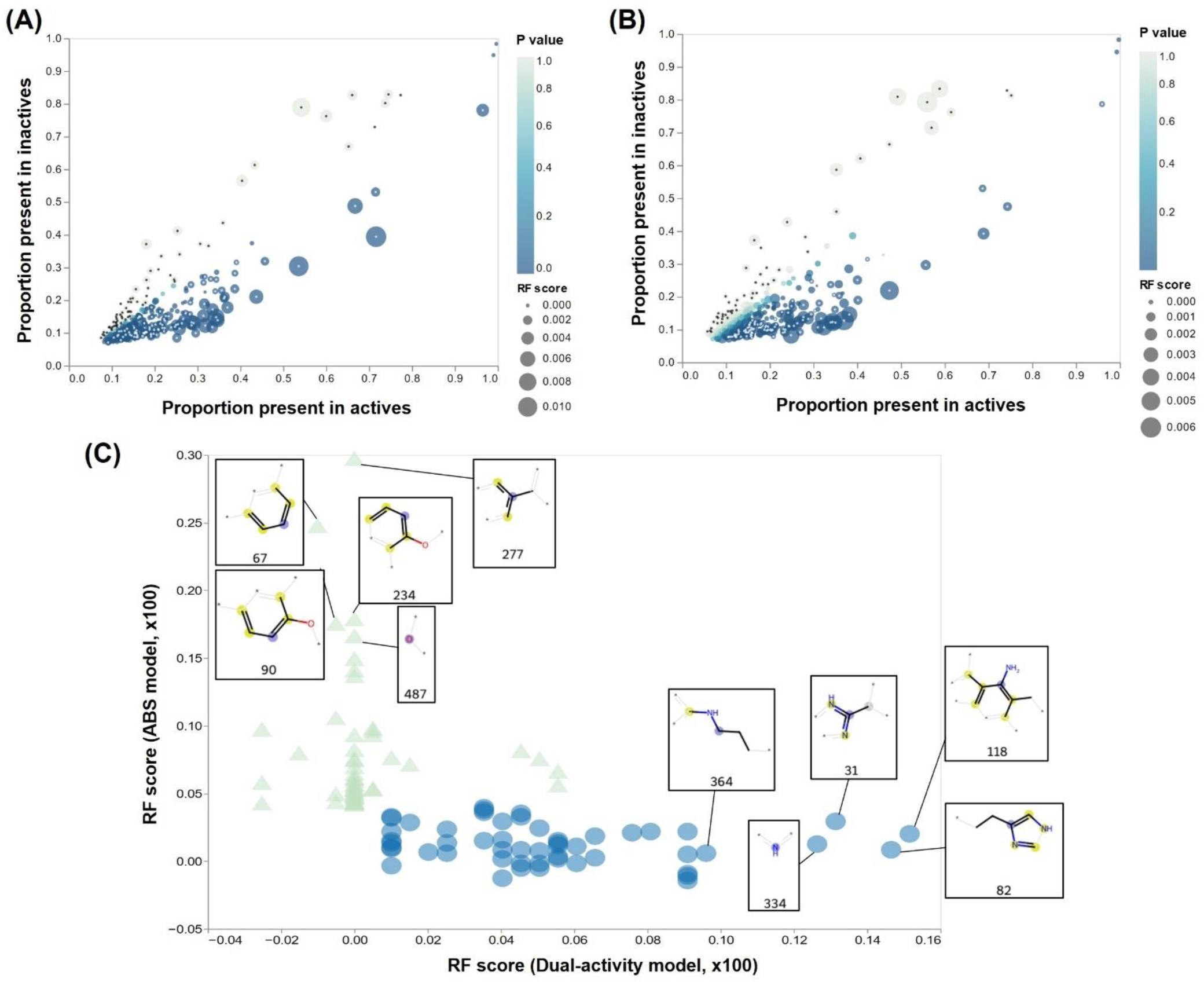
Enriched Mfp features within inactive and active compounds for stage-specific antiplasmodial action. **(A)** The proportion of active/inactive compounds against ABS containing a specific Mfp feature is plotted as circles, with the size of the circles corresponding to the RF permutation score of the Mfp feature. Enrichment of a feature towards active compounds compared to inactive compounds is indicated by the p-value color obtained from the Z-test on two proportions. The top 100 enriched Mfp features within active (white) and the top 67 enriched Mfp features within inactive (black) compounds were selected according to RF score and ρ-value. **(B)** The proportion of dual-active/inactive compounds containing a specific Mfp feature is plotted as circles, with the size corresponding to the RF permutation score of the Mfp feature. Enrichment of a feature towards dual-active compounds compared to inactive compounds is indicated by the ρ-value color. The top 100 enriched Mfp features within dual-active (white) and the top 52 enriched Mfp features within inactive (black) compounds were selected according to RF score and ρ-value. **(C)** Comparison of the unique Mfp features associated with activity against ABS (52) or dual stages (52). For the top unique Mfp features, structural elements are indicated with all features summarised in Supplementary file 2.

As expected, around 50% of the top 100 features associated with activity were shared between compounds having ABS inhibition activity or dual activity. However, importantly, of the top 100 features enriched in the respective active fractions, 52 features each uniquely described features either associated with sole ABS activity or with dual-activity (Figure 4C, Supplementary file S2). This provides clear distinction between chemical composition of compounds required to kill multiple stages of the parasite compared to that required to only kill ABS parasites. Indeed, the top 5 unique features enriched in compounds able to target ABS parasites contain heterocyclic structures that are activated via oxygenation or alkylation, making the compound more reactive to electrophilic attack. Oxygenation may relate to a compound’s bioavailability by increasing the compound’s solubility and uptake ^39^. Comparatively, the top 5 unique features that are enriched in compounds with dual-stage activity are enriched for amine groups whether it be nitrogen containing 4-membered heterocyclic structures or the nitration of benzene and/or carbon groups. Such amine groups typically create localized electron deficient sites in the compound that allows interaction with cellular components such as amino acids and nucleic acids to be more favourable ^40^. Hence, such amine groups may be more involved in the killing/drug effect as well as aid in solubility of compounds.

Together with this, chemical features enriched in compounds that are inactive against ABS parasites (67) and/or gametocytes (52) tended to overlap with one another. Generally, these features indicate that features where nitrogen is unable to bind to hydrogen or structures containing amides and branching carbon chains may be associated with a lack in bioactivity (Supplementary file S2).

### 3.5. Model validation in novel chemical spaces

To further validate and interrogate the extent in which the above models can predict stage-specific activity of antimalarial compounds, the SVM and RF models were exposed to previously unseen chemical matter and data from curated datasets. These datasets served as an added level of interrogation to determine how well our models would perform when exposed to very diverse and novel chemical spaces and can give an indication of what to expect of these models in real world application. We used data from the MMV PRB and Pathogen box, which contained unique compounds not included in the data from the chemical libraries used to train the models, and with activity data available against both *P. falciparum* ABS parasites and gametocytes ^3, 26^. The compounds included in the boxes were also individually distinct and chemically diverse, providing more extreme datasets to evaluate the robust nature of the models, compared to the larger databases used to train the models, where structurally related compounds were present within a chemical space (Figure 5A and D). The best model must thus maintain fair accuracy and recall under these conditions, whilst optimising sensitivity and specificity in predictions to limit the models’ FPR for these novel and diverse chemical matter. Class imbalance was not corrected for these datasets to evaluate the performance of the models, with hit rates against ABS at 18% for the PRB box ^3^ and 31% for the Pathogen Box ^26^ (Figure 5A and D). For compounds with gametocytocidal activity the hit rate for the PRB box (∼13%) and Pathogen box (∼24%) was even lower.

**Figure 5.**
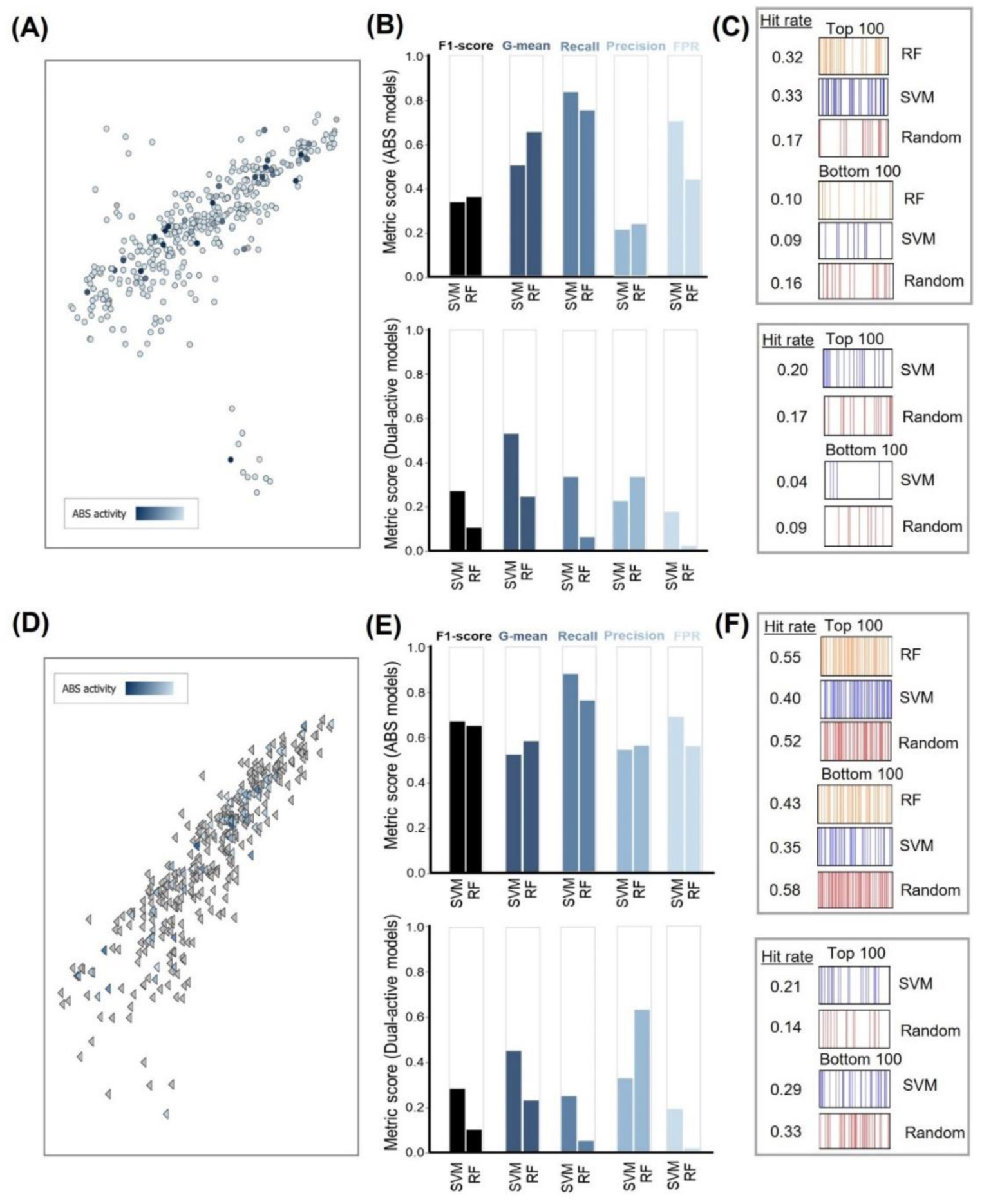
Performance of the top models against unseen chemical matter. To evaluate model robustness, models were exposed to extreme datasets, such as the PRB box and Pathogen box that were individually distinct, chemically diverse, and had differential activity against ABS and gametocytes. **(A)** and **(D)** indicate the diversity of the compounds included in these sets, displayed within context of the launched drug chemical space (available on StarDrop v 7.3.0), with heatbars indicating increased potency. ABS and dual-activity models trained on Mfp descriptors of compounds were evaluated for their activity predictions within the PRB box **(B)** and the Pathogen box **(E)**. F1-score evaluated how model performed when exposed to imbalanced data, whereas G-mean scores how well models were able to optimise sensitivity and specificity. Recall and precision indicated accuracy of activity predictions whereas false positive rate (FPR) indicated error within predictions. The hit rate of the best performing model for predicting ABS and/or dual-activity within these chemical spaces **(C and F)** were compared to random selection. Models ranked compounds according to probability of having activity and the top and bottom 100 compounds from the models and random selection were used to calculate the hit rate with the ideal having a high top 100 hit rate and low bottom 100 hit rate compared to random selection.

Within the two top-performing ABS activity models (SVM and RF), both models obtained similar scores for all metrics on the PRB box (Figure 5B). Both models were able to show a significant enrichment of hits (hit rates at >30%) in the top 100 compounds, compared to random selection (17%) and the hit rate of the PRB box itself (18%) (Figure 5C). The specificity is also confirmed as these models enable selection of compounds that would be inactive in the bottom 100. Within this chemical space, the models differentiated from each other when predicting activity for dual-active compounds, with the SVM model outperforming on all metrics, excluding precision (Figure 5B). Although SVM tended to have a higher FPR (0.17 vs 0.02), RF failed to identify dual-active compounds with a recall below <0.20. Together with this the hit rate of the top 100 hits from the SVM model (20%) exceeded the hit rate of random selection (17%) and the PRB box (13%) (Figure 5C).

Within a different novel chemical space (Pathogen Box, Figure 5D), both SVM and RF ABS activity models obtained higher F1-scores (Figure 5E), compared to the PRB box (Figure 5B) and this can be attributed to the higher hit rate within the Pathogen Box resulting in a less severe class imbalance compared to the PRB box. Model performance was maintained (Figure 5E) as also evident in enrichment of hits (>40%) for predicting ABS activity as well as dual active compounds (Figure 5F) With the Pathogen Box, the dual-activity models maintained similar F1- scores when predicting dual-active compounds, indicating that despite the difference in hit rates between the PRB and Pathogen box, the dual-active models are not influenced by class imbalance, which is highly beneficial. Although the dual-activity models’ performance is lower compared to the test set, this was expected considering the diverse data the models were exposed to and the limited chemical space data available for training in comparison to the ABS database.

## 4. DISCUSSION

The identification of gametocytocidal compounds remains challenging compared to identifying compounds with activity against ABS parasites, requiring alternative approaches to streamline screening and compound optimisation campaigns. To our knowledge, no study has fully explored the chemical space information of phenotypic screens conducted against gametocytes to help in predicting gametocytocidal compounds, although this has been done for asexual phenotypic screens via DeepMalaria and MAIB ^18, 20^ as well as liver-stage phenotypic screens ^19^. Here, we were able to successfully train models capable of predicting compounds with activity against both ABS parasites and gametocytes with high accuracy, precision and recall.

Our best performing SVM and RF models for ABS inhibition activity prediction were on par with DeepMalaria ^18^ that used neural network based models, with SVM obtaining higher recall abilities (95% *vs*. 87.75%) but slightly less precision (2.40% *vs*. 3.54%) on untrained data. This could be because the chemical space information we used for training was larger as we incorporated more chemical libraries screened against ABS. Additionally, the cluster-based undersampling of inactive compounds as used here seems to be a more important strategy compared to the oversampling of active compounds used before. One disadvantage of oversampling can be that the model fails to generalize and recognize patterns in novel active compounds due to the model being fixated on patterns associated with the oversampled active compounds as a result of such features being over represented in the training data ^38^. Our data therefore suggest that more simplified algorithms, such as those used here, together with strict attention to class bias, could be more informative in building ML models for the antimalarial chemical space, negating the use of more complex approaches using neural network analysis. Similar observations have been seen in different fields involving image classification when comparing deep learning to simpler machine learning methods such as SVM ^41^. Single classifiers such as SVM outperformed ensemble models within representative chemical spaces in our data, which could be considered smaller than the typical big data on which ensemble methods such as GBM and RF generally performs better. Alternatively, the fact that SVM algorithms balances the contributions of chemical features in correct predictions could be more important in the antimalarial drug space compared to the way ensemble methods rely mostly only on the presence/absence of such features ^42^.

Within this study, we identified chemical features enriched within dual-active compounds. Interestingly, Mfp molecular featurization enabled models to perform better within novel and diverse antimalarial chemical spaces compared to MACCS keys. This could be attributed to Mfp capturing the atom environments around each atom, whereas MACCS summarises the presence/absence of chemical groups within compounds that have been identified as important within drug discovery and virtual screening ^43^ with both these featurizations incorporated into tools such as Chemical Checker ^44^. Hence, Mfp due to them being more descriptive may be better at capturing the chemical space and translating this over to models, whereby such models can then identify chemical features important for bioactivity as well as a lack in bioactivity. The most notable commonality amongst enriched features was the nitration of heterocyclic and carbon groups that may aid in interactions with cellular targets ^40^. Similarly for features enriched within ABS inhibiting compounds and important for predicting ABS inhibition, oxygenation was more common and may relate to solubility and drug uptake. This study therefore contributes the identification for chemical features important for stage-specific activity and we anticipate that inclusion of chemical features (or exclusion of those features associated with inactivity) will aid medicinal chemist to guide compound derivatization during hit-to-lead optimisation of stage-specific and/or dual active candidates. However, it must be noted that the features described here lack the context of connectivity and it is more than likely that a combination of such features is important for stage-specific activity. The inclusion of additional molecular and biological descriptors with physicochemical properties ^44^ may enable us to better understand how chemical features contribute to stage-specific activity of compounds and possibly deconvoluting mode of action. Alternatively, the capabilities of generative models may be useful to artificially expand this limited chemical space by creating *de novo* chemical starting points with a high probability of having desirable gametocytocidal activity and drug-like properties ^45^.

## 5. CONCLUSIONS

In summary, this study contributes a new ML tool to guide and optimise both phenotypic screens as well as compound derivatisation in hit-to-lead and lead optimisation campaigns, so as to reduce time and cost allocated towards compounds with low probability of having dual activity against both ABS parasites and gametocytes. Considering such models have good understanding of the chemical space for bioactivity, we were successful in mining for features related to bioactivity and highlight chemical features predictive towards ABS and/or dual-activity. The tools provided by this study can accelerate the identification and hit-to-lead optimisation of dual-active compounds to aid in malaria elimination strategies. These models are deployed within the Ersilia Model Hub repository ^46^ for open-source access, particularly aiding infectious disease drug discovery.

## DATA AVAILABILITY STATEMENT

The datasets used in this study can be found at ChEMBL using ChEMBL ID provided in the Supplementary file S1 or through accessing the original articles cited.

## CODE AVAILABILITY STATEMENT

All python scripts for clustering, undersampling and model building as well as evaluation can be obtained from github: <u>github.com/M2PL/Machines-Against-Malaria</u>. To facilitate model usage, we have also incorporated the models in the Ersilia Model Hub (https://www.ersilia.io/model-hub; identifier eos80ch).

## ASSOCIATED CONTENT

**Supporting Information**. The following files are available free of charge.

**Supplementary document** contains information on hyperparameters used for model building and additional performance metrics of model predictions in representative and novel chemical spaces. (DOC)

**Supplementary file S1** contains chemical information of compounds within the databases used for ML as well as performance metrics of models trained on imbalanced/oversampled/undersampled data using either Mfp or MACCS molecular fingerprints. (XLSX)

**Supplementary file S2** contains information of enriched Mfp features for activity/inactivity against ABS and/or gametocytes. (XLSX)

**SMILES** contains Simplified Molecular Input Line Entry System (SMILES) of compounds used for machine learning. (CSV)

### Author Contributions

AvH performed the research with inputs from MDF, GT and NP. LMB conceptualized the study and wrote the paper with AvH. All co-authors approved the final version of the paper.

### Funding Sources

This work was supported by the South African Department of Science and Innovation and National Research Foundation South African Research Chairs Initiative Grant (LMB UID: 84627).

### Notes

The authors declare that the research was conducted in the absence of any commercial or financial relationships that could be construed as a potential conflict of interest.

## ACKNOWLEDGMENT

We thank Jason Hlozeck from the University of Cape Town for useful discussions and proofreading of the paper. Additionally, the authors acknowledge Google Colab for their cloud-based service that allowed us to build machine learning models.

## ABBREVIATIONS

ABS: Asexual blood stages
ACT: artemisinin combination therapy
FPR: false positivity rate
GBM: Gradient Boosting Machines
G-mean: geometric mean
LR: Logistic Regression
MACCS: Molecular Access keys
Mfp: Morgan fingerprint
ML: Machine learning
PRB: Pandemic Response Box
SVM: Support Vector Machines
RBF: radial basis function
RF: Random Forests
ROC-AUC: receiver operator characteristic curve
TrB: transmission-blocking.

