## Supplementary document for "Machine learning approaches identify chemical features for stage-specific antimalarial compounds"

### Supplementary Material

The optimal bit-length doe ABS activity prediction models was determined to be 500-bit as there appeared to be no change in ROC-AUC values as bit length was increased, whereas at a drastic drop in FPR was observed at 500-bit which then again increased as the bit-length was increased. In contrast to FPR, the opposite trend was observed for precision. For Dual activity prediction models, ROC-AUC values slightly increased as bit length increased with GBM decreasing in ROC-AUC once larger than 500-bit length was used. FPR values decreased with increasing bit length, however models plateaued at 300-bit length (excluding LR). Precision also tended to increase with increasing bit-length however, at 500-bits only slight increases was observed at 500-bits. Based on ROC-AUC, FPR and precision metrics as well as the training time required to build models 500-bits were determined to be the best bit length to use for Mfp whilst training most models.

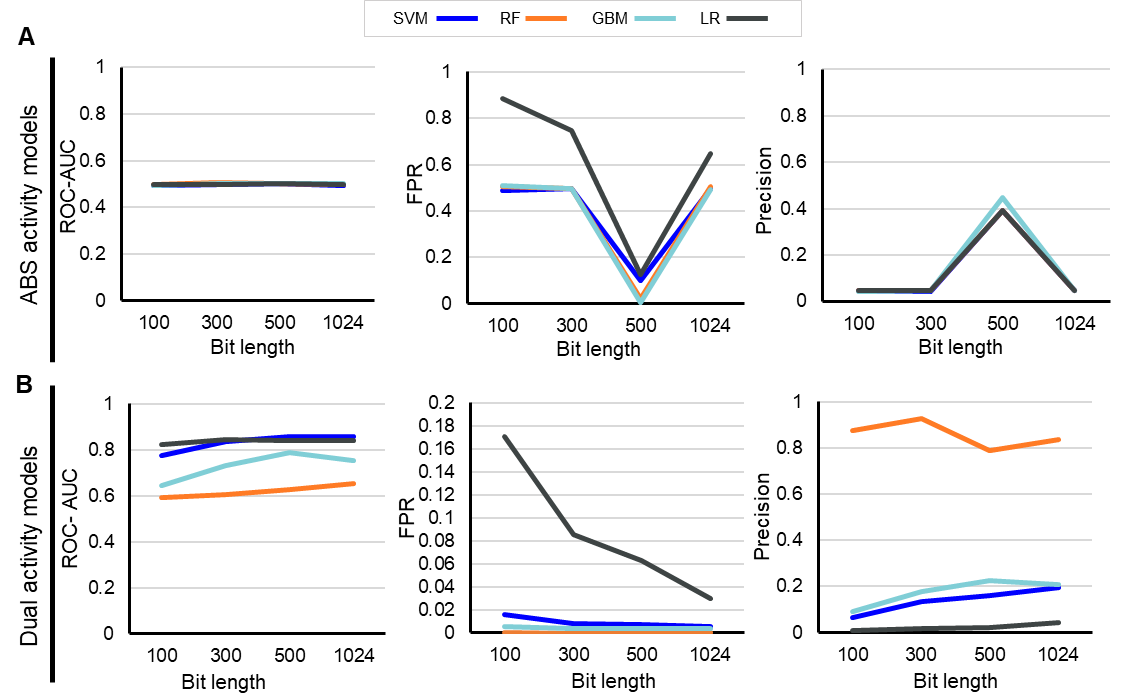

**Supplementary Figure S1: Optimal Morgan fingerprints (Mfp) bit length that enabled better model performance.**

(A) Performance of ABS activity models using different ML algorithms (blue = Support vector machine, orange = Random forest, light blue= gradient boosting machine, dark grey= logistic regression) in identifying compounds with ABS inhibition activity within test set using differing bit lengths of Morgan fingerprints (Mfp) during training. ROC AUC scores indicate the classifier’s ability in distinguishing active and inactive compounds against ABS. FPR scores indicate false positive rate of test set predictions. (B) Performance of dual active models (trained on data with class imbalance) using different ML algorithm’s (blue = Support vector machine, orange = Random forest, light blue= gradient boosting machine, dark grey= logistic regression) ability in identifying compounds with dual activity within test set using differing bit lengths of Morgan fingerprints (Mfp) during training.

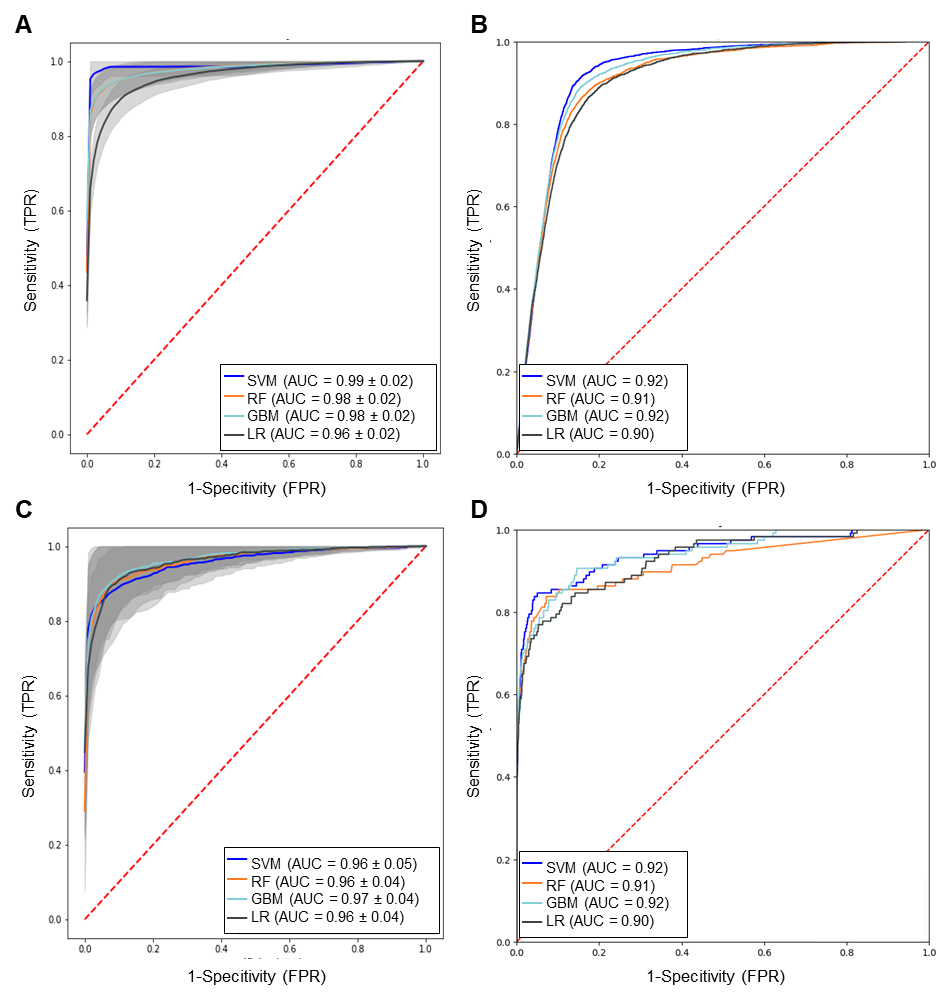

**Supplementary Figure S2: ROC-AUC curves of ABS and dual-activity MACCS models on 5-fold cross-validation and untrained test set.**

ROC-AUC curves showing performance of different ML algorithms in predicting compounds with ABS (A) or (C) dual-activity when trained on MACCS keys of compounds after 5-fold cross-validation. Insert indicates AUC mean values ± standard deviation. The ROC-AUC curves of the different models trained on MACCS descriptors on untrained test set in predicting ABS (B) or dual-activity (D).

**Table S1: Hyperparameter tuning and optimal parameters identified for models**

| Parameter tuned | Parameters (range) used | Optimal parameter (Mfp) | Optimal parameter  (MACCS) |
| --- | --- | --- | --- |
| Dual activity prediction SVM model | | |  |
| Kernels: | Polynomial, RBF, Sigmoid, Linear | RFB | RFB |
| Regularization parameter (C) | [0.1, 1, 10, 100, 1000] or default | 10 | 1000 |
| Kernel coefficient (gamma) | [1, 0.1, 0.01, 0.001, 0.0001] or default | 0.01 | 0.01 |
| ABS activity prediction SVM model | | |  |
| Kernels: | Polynomial, RBF, Sigmoid, Linear | RFB | RFB |
| Regularization parameter (C) | [0.1, 1, 10, 100, 1000] or default | default | 10 |
| Kernel coefficient (gamma) | [1, 0.1, 0.01, 0.001, 0.0001] or default | default | 0.1 |
| GBM dual activity prediction model | | |  |
| Number of trees | [10, 50, 100, 500] | 500 | 500 |
| Subsample | [0.1, 0.01, 0.001] | 0.1 | 0.1 |
| Max features | 1-10 | 5 | - |
| Learning rate | [1, 0.1, 0.01, 0.001, 0.0001] | 0.1 | 0.01 |
| Max tree depth | 1-10 | 2 | 9 |
| ABS activity prediction GBM model | | |  |
| Number of trees | [10, 50, 100, 500] | 100 | 500 |
| Subsample | [0.1, 0.01, 0.001] | 0.1 | 0.1 |
| Max features | 1-7 | 29 | - |
| Learning rate | [1, 0.1, 0.01, 0.001, 0.0001] | 0.1 | 0.01 |
| Max tree depth | 1-10 | 2 | 9 |
| RF dual activity prediction model | | |  |
| Max depth | [10- 15] | 14 | 14 |
| Max features | ['auto', 'log2'] | auto | auto |
| Number of estimators | [5, 6, 7, 8, 9, 10, 11, 12, 13, 15] | 15 | 10 |
| ABS RF activity prediction model | | |  |
| Max depth | 10-15 | 14 | 14 |
| Max features | ['auto', 'log2'] | auto | auto |
| Number of estimators | [5, 6, 7, 8, 9, 10, 11, 12, 13, 15] | 15 | 15 |
| LR dual activity prediction model | | |  |
| C-value | [100, 10, 1.0, 0.1, 0.01] | 100 | 100 |
| Solvers | ['newton-cg', 'lbfgs', 'liblinear'] | lbfgs | lbfgs |
| Penalty | L2 | L2 | L2 |
| ABS LR activity prediction model | | |  |
| C-value | [100, 10, 1.0, 0.1, 0.01] | 10 | 100 |
| Solvers | ['newton-cg', 'lbfgs', 'liblinear'] | lbfgs | newton-cg |
| Penalty | L2 | L2 | L2 |

**Table S2: Optimised probability threshold**

| Model | G-Mean | FPR | ROC-AUC | Recall | Precision | Probability threshold |
| --- | --- | --- | --- | --- | --- | --- |
| ABS activity prediction model trained on undersampled balanced data | | | | | | |
| SVM (Mfp) | 0.875 | 0.149 | 0.875 | 0.899 | 0.224 | 0.5 |
| RF (Mfp) | 0.825 | 0.178 | 0.825 | 0.828 | 0.182 | 0.5 |
| GBM (Mfp) | 0.802 | 0.219 | 0.802 | 0.824 | 0.152 | 0.5 |
| LR (Mfp) | 0.708 | 0.483 | 0.743 | 0.969 | 0.088 | 0.5 |
| SVM (Mfp) | 0.714 | 0.468 | 0.745 | 0.958 | 0.675 | 0.8 |
| RF (Mfp) | 0.702 | 0.415 | 0.714 | 0.842 | 0.673 | 0.64 |
| GBM (Mfp) | 0.706 | 0.071 | 0.733 | 0.537 | 0.267 | 0.74 |
| LR (Mfp) | 0.822 | 0.140 | 0.823 | 0.785 | 0.212 | 0.96 |
| Dual activity prediction model trained on undersampled balanced data | | | | | | |
| SVM class-weighted (Mfp) | 0.809 | 0.006 | 0.826 | 0.658 | 0.164 | 0.5 |
| RF (Mfp) | 0.562 | 0.000 | 0.658 | 0.316 | 0.578 | 0.5 |
| GBM (Mfp) | 0.720 | 0.005 | 0.758 | 0.521 | 0.159 | 0.5 |
| LR class-weighted (Mfp) | 0.796 | 0.061 | 0.807 | 0.675 | 0.020 | 0.5 |
| SVM class-weighted (Mfp) | 0.654 | 0.001 | 0.713 | 0.427 | 0.562 | 0.92 |
| RF (Mfp) | 0.562 | 0.000 | 0.658 | 0.316 | 0.617 | 0.52 |
| GBM (Mfp) | 0.547 | 0.000 | 0.649 | 0.299 | 0.565 | 0.82 |
| LR class-weighted (Mfp) | 0.768 | 0.041 | 0.787 | 0.615 | 0.027 | 0.96 |

**Table S3: Complex model Autogluon comparison on test data**

| Model | G-Mean | FPR (FP/FP+TN) | ROC-AUC | Recall | Precision | F1-Score |
| --- | --- | --- | --- | --- | --- | --- |
| ABS activity prediction model | | | | | | |
| SVM (Mfp) | 0.875 | 0.149 | 0.875 | 0.899 | 0.224 | 0.359 |
| RF (Mfp) | 0.825 | 0.178 | 0.825 | 0.828 | 0.182 | 0.298 |
| NeuralNetFastAI | 0.863 | 0.188 | 0.864 | 0.917 | 0.189 | 0.313 |
| WeightedEnsemble_L2 | 0.877 | 0.163 | 0.877 | 0.918 | 0.213 | 0.345 |
| Dual activity prediction model | | | | | | |
| SVM (Mfp) | 0.809 | 0.006 | 0.826 | 0.658 | 0.164 | 0.485 |
| RF (Mfp) | 0.562 | 0.000 | 0.658 | 0.316 | 0.578 | 0.409 |
| NeuralNetFastAI | 0.992 | 0.632 | 0.813 | 0.632 | 0.147 | 0.238 |
| WeightedEnsemble_L2 | 0.995 | 0.641 | 0.818 | 0.641 | 0.201 | 0.305 |

**Table S4: Metrics per model on PRB and Pathogen box data**

| Model | G-Mean | FPR (FP/FP+TN) | Sensitivity (TP/TP+FN) | Specificity (TN/TN+FP) | F1-Score |
| --- | --- | --- | --- | --- | --- |
| ABS inhibition activity prediction model on PRB box | | | | | |
| SVM (Mfp) | 0.499 | 0.701 | 0.833 | 0.299 | 0.331 |
| RF (Mfp) | 0.650 | 0.437 | 0.750 | 0.563 | 0.356 |
| GBM (Mfp) | 0.547 | 0.585 | 0.722 | 0.415 | 0.329 |
| LR (Mfp) | 0.327 | 0.890 | 0.972 | 0.110 | 0.323 |
| SVM (MACCS) | 0.432 | 0.784 | 0.861 | 0.216 | 0.317 |
| RF (MACCS) | 0.446 | 0.748 | 0.792 | 0.252 | 0.302 |
| GBM (MACCS) | 0.007 | 0.999 | 0.059 | 0.001 | 0.313 |
| LR (MACCS) | 0.255 | 0.933 | 0.972 | 0.067 | 0.313 |
| ABS inhibition activity prediction model on Pathogen box | | | | | |
| SVM (Mfp) | 0.521 | 0.689 | 0.876 | 0.311 | 0.670 |
| RF (Mfp) | 0.581 | 0.558 | 0.763 | 0.442 | 0.646 |
| GBM (Mfp) | 0.536 | 0.600 | 0.718 | 0.400 | 0.608 |
| LR (Mfp) | 0.215 | 0.953 | 0.977 | 0.047 | 0.652 |
| SVM (MACCS) | 0.493 | 0.726 | 0.887 | 0.274 | 0.665 |
| RF (MACCS) | 0.532 | 0.626 | 0.757 | 0.374 | 0.623 |
| GBM (MACCS) | 0.511 | 0.663 | 0.774 | 0.337 | 0.623 |
| LR (MACCS) | 0.296 | 0.911 | 0.977 | 0.089 | 0.662 |
| Dual activity prediction model on PRB box | | | | | |
| SVM class-weighted (Mfp) | 0.525 | 0.172 | 0.333 | 0.828 | 0.266 |
| RF (Mfp) | 0.240 | 0.017 | 0.059 | 0.983 | 0.100 |
| GBM (Mfp) | 0.487 | 0.135 | 0.275 | 0.865 | 0.250 |
| LR class-weighted (Mfp) | 0.593 | 0.309 | 0.510 | 0.691 | 0.281 |
| SVM class-weighted (MACCS) | 0.521 | 0.341 | 0.412 | 0.659 | 0.220 |
| RF (MACCS) | 0.444 | 0.160 | 0.235 | 0.840 | 0.202 |
| GBM (MACCS) | 0.501 | 0.246 | 0.333 | 0.754 | 0.221 |
| LR class-weighted (MACCS) | 0.571 | 0.550 | 0.725 | 0.450 | 0.264 |
| Dual activity prediction model on Pathogen box | | | | | |
| SVM class-weighted (Mfp) | 0.449 | 0.185 | 0.247 | 0.815 | 0.281 |
| RF (Mfp) | 0.226 | 0.011 | 0.052 | 0.989 | 0.095 |
| GBM (Mfp) | 0.444 | 0.170 | 0.237 | 0.830 | 0.277 |
| LR class-weighted (Mfp) | 0.467 | 0.359 | 0.340 | 0.641 | 0.291 |
| SVM class-weighted (MACCS) | 0.492 | 0.289 | 0.340 | 0.711 | 0.317 |
| RF (MACCS) | 0.408 | 0.104 | 0.186 | 0.896 | 0.252 |
| GBM (MACCS) | 0.408 | 0.152 | 0.196 | 0.848 | 0.242 |
| LR class-weighted (MACCS) | 0.567 | 0.463 | 0.598 | 0.537 | 0.414 |
